# Agreement between ^18^F-Florbetapir PET imaging and cerebrospinal fluid Aβ1-42, Aβ1-40, tTau and pTau measured on the LUMIPULSE G fully automated platform

**DOI:** 10.1101/476937

**Authors:** Daniel Alcolea, Jordi Pegueroles, Laia Muñoz, Valle Camacho, Diego López-Mora, Alejandro Fernández-León, Nathalie Le Bastard, Els Huyck, Alicia Nadal, Verónica Olmedo, Victor Montal, Eduard Vilaplana, Jordi Clarimón, Rafael Blesa, Juan Fortea, Alberto Lleó

## Abstract

**INTRODUCTION:** The development of fully automated immunoassay platforms has improved the technical reliability of cerebrospinal fluid (CSF) biomarkers for Alzheimer’s disease.

**METHODS:** We quantified Aβ1-42, Aβ1-40, tTau and pTau levels using the Lumipulse G System in 94 CSF samples from participants of the SPIN cohort with available ^18^F-Florbetapir imaging. Amyloid scans were assessed visually and through automated quantification. We determined the cutoffs of CSF biomarkers that optimized their agreement with ^18^F-Florbetapir PET and evaluated concordance between markers of the amyloid category.

**RESULTS:** Aβ1-42, tTau and pTau (but not Aβ1-40) and the ratios with Aβ1-42 had good diagnostic agreement with ^18^F-Florbetapir PET. As a marker of amyloid pathology, the Aβ1-42/Aβ1-40 ratio had higher agreement and better correlation with amyloid PET than Aβ1-42 alone.

**DISCUSSION:** CSF biomarkers measured with the Lumipulse G System show good agreement with amyloid imaging. Combination of Aβ1-42 with Aβ1-40 increases the agreement between markers of amyloid pathology.

## 1. Introduction

Advances in the field of biomarkers have pushed forward a redefinition of Alzheimer’s disease (AD) as a biological construct [1]. Under this new definition, the ATN classification system recognizes three general groups of biomarkers for AD based on the pathologic process that they reflect: biomarkers of β-amyloid plaques (A), biomarkers of fibrillar tau (T) and biomarkers of neurodegeneration or neuronal injury (N) [1]. These biomarkers have been incorporated in clinical trials for patients’ selection and to monitor target engagement [1–3]. In clinical practice, biomarkers are useful to detect or exclude AD, to make a prognosis, and to guide patients’ management, particularly in atypical and clinically challenging cases [4].

Biomarkers of different modalities have been investigated in AD, but those that are more widely implemented are cerebrospinal fluid (CSF) biomarkers and imaging techniques. In CSF, a combination of low levels of Aβ1-42 and high levels of total tau (tTau) and 181-phosphorylated tau (pTau) has proven high accuracy to detect AD pathophysiology, even before the appearance of symptoms [5]. Pathological studies have shown that low levels of Aβ1-42 in CSF are associated to high amyloid plaques deposition [6,7], and levels of tTau and pTau in CSF correlate with neuronal loss and neurofibrillary tangle burden [7,8]. In the molecular imaging field, amyloid and Tau tracers provide *in vivo* topographical knowledge of the amount and distribution of pathology. The ATN classification system groups amyloid PET, CSF Aβ1-42 and the ratio Aβ1-42/Aβ1-40 in the “A” category, Tau PET and CSF pTau in the “T” category, whereas ^18^F-Fluorodeoxyglucose-PET and tTau are considered markers of the “N” category [1]. CSF biomarkers are thus very informative and relatively inexpensive, but obtaining them requires a lumbar puncture, and they do not reveal the distribution of pathology. The use of molecular imaging is less invasive but, in turn, radiotracers are more expensive, radioactive, and only accessible in centers with the appropriate facilities. The choice of whether to use CSF or PET biomarkers will depend on a variety of factors, including availability, patient comorbidities and cost.

The concordance between low CSF Aβ1-42 levels and positive amyloid PET imaging is high, but not perfect [9–13]. One of the reasons that may explain discordant results between these two modalities is that, while amyloid imaging reflects the progressive accumulation of brain fibrillar amyloid deposits, decreases in soluble Aβ1-42 levels in CSF can be attributed to pathological processes other than amyloid plaque accumulation [14]. Aβ1-42 peptide is the result of a sequential proteolytic cleavage of the amyloid precursor protein, and its soluble levels in CSF might be affected by individual differences in the abundance of amyloid precursor protein or in the rate of its cleavage [15,16]. Previous studies have shown that the combination of Aβ1-42 with other isoforms of Aβ such as Aβ1-40 (Aβ1-42/Aβ1-40) can correct these individual differences. In fact, the Aβ1-42/Aβ1-40 ratio has shown to have better agreement with amyloid PET imaging compared to levels of Aβ1-42 alone [11,14,17,18] and to be a key marker in clinical practice for the evaluation of AD [15]. Another factor that might contribute to discordances between amyloid imaging and CSF biomarkers is the fact that traditional ELISA assays have large inter-assay variability ranging from 20% to 30% [19]. In recent years, the development of fully automated platforms for the analysis of CSF biomarkers has reduced this variability to less than 10% [20,21]. The development of technically reliable platforms is a crucial step for their implementation in clinical routine, for the calculation of accurate cutoffs and for studying the association of CSF biomarkers with markers of other modalities. Recently, four CSF analytes (Aβ1-42, Aβ1-40, tTau and pTau) have been implemented on the fully automated Lumipulse G System, but there are, however, no validated cutoffs for these four AD CSF biomarkers using this platform. Our aims were to determine for the first time the cutoffs that optimized the agreement between ^18^F-Florbetapir PET and Aβ1-42, Aβ1-40, tTau, pTau and their ratios measured in CSF on the LUMIPULSE G600II instrument, and to evaluate the concordance between markers of the amyloid category.

## 2. Methods

### 2.1 Study participants

A set of participants of the Sant Pau Initiative on Neurodegeneration (SPIN cohort) was included in the study. The SPIN cohort is a multimodal research cohort for biomarker discovery and validation that includes participants with different neurodegenerative dementias, mild cognitive impairment and cognitively normal controls. All participants receive an extensive neurological and neuropsychological evaluation and undergo structural 3T brain MRI, blood extraction, and lumbar puncture for CSF biomarkers. A subset of participants also receives molecular imaging such as ^18^F-Fluorodeoxyglucose-PET, amyloid and/or Tau PET. More information on the SPIN cohort can be found at https://santpaumemoryunit.com/our-research/spin-cohort (Alcolea et al., in preparation).

In this study, we included all 94 participants from the SPIN cohort recruited between November 2013 and September 2017 that had available CSF samples and ^18^F-Florbetapir PET imaging. Their clinical diagnoses were mild cognitive impairment (n=35), AD dementia (n=12), other dementias or neurodegenerative diseases (that included dementia with Lewy bodies [n=30], frontotemporal dementia [n=9], vascular dementia [n=1], and motor neuron disease [n=1]), and cognitively normal controls (n=6). All participants gave written consent, and the ethics committee of Hospital Sant Pau approved all procedures included in this study.

### 2.2 CSF samples acquisition and analysis

CSF samples were collected in 10 ml polypropylene tubes (Sarstedt, Ref# 62.610.018) and immediately transferred to the Sant Pau Memory Unit’s laboratory where they were processed within the first 2 hours after acquisition. After centrifugation (2000 g x 10 min, 4ºC), volumes of 0.5 ml of CSF were aliquoted into polypropylene tubes (Sarstedt, Ref# 72.694.007) and stored at −80ºC until analysis.

On the day of the analysis, samples were thawed at room temperature and the tubes were vortexed for 5-10 seconds. To avoid the effect of multiple freeze-thaw cycles, aliquots used in this study had not been thawed before. Aβ1-42, Aβ1-40, tTau and pTau were quantified directly from the storage tubes containing 0.5 ml of CSF using the Lumipulse G β-Amyloid 1-42, β-Amyloid 1-40, Total Tau and pTau 181 assays by the LUMIPULSE G600II automated platform and following the manufacturer’s instructions. We used the same batch of reagents for each biomarker throughout all the study, and for each sample, we measured all four analytes from the same aliquot and in the same run. The platform was configured to start the analysis with Aβ1-42, followed by Aβ1-40, tTau and pTau. Quality control testing was performed at the beginning of each test day to ensure that all measured values of each control level (low, medium and high) were within the target ranges.

The results of the Lumipulse G β-Amyloid 1-42 presented in this study have been standardized according to certified reference material developed by the International Federation of Clinical Chemistry and Laboratory Medicine as recommended by their working group for CSF proteins [22]. Briefly, values of the calibration standards of the LUMIPULSE G600II were adapted to the certified reference material resulting in an adjustment of concentrations that was linearly proportional throughout all the range. The aim of standardization to certified reference material is to harmonize immunoassays of Aβ1-42 to make results comparable across different platforms.

Aβ1-42, Aβ1-40, tTau and pTau levels in CSF from participants of this study had been measured previously using other immunoassays (INNOTEST β-AMYLOID_(1-42)_, INNOTEST hTAU Ag, and INNOTEST PHOSPHO-TAU_(181P)_, Fujirebio Europe; and High Sensitivity Human Amyloid β40, Merck-Millipore), and these results were available in our database for their comparison with the LUMIPULSE analyses [23–27]. The personnel involved in the CSF analyses for this study were blinded to the clinical diagnosis and to previous biomarker determinations.

### 2.3 Amyloid-PET imaging acquisition, visual assessment and quantitative analysis

All participants underwent amyloid PET imaging with ^18^F-Florbetapir as described elsewhere [25]. PET data were acquired using a Philips Gemini TF scan 50 minutes after injection of 370 mBq of ^18^F-Florbetapir. After obtaining the transmission data, brain PET dynamic acquisition was performed (2 x 5 min frames). The reconstruction method was iterative (LOR RAMBLA, 3 iterations and 33 subsets) with a 128 x 128 image size, 2 mm pixel size and slice thickness.

Three expert readers (V.C., D.L-M. and A.F-L.) that were blind to clinical diagnosis and to CSF biomarker results visually rated all PET scans. Following manufacturer’s protocol, scans were classified as “positive” when one or more areas showed increased cortical gray matter signal resulting in reduced or absent contrast between gray matter and white matter. Scans were classified as “negative” when the contrast between gray matter and white matter was clear. Final classification as “positive” or “negative” was decided upon agreement of at least two of three readers. Mean inter-reader overall agreement was 88.4% (Min=87.0%, Max=90.2%).

We also quantified amyloid deposition. Briefly, each participant's PET scan was spatially normalized to a MNI152 ^18^F-Florbetapir template using a linear and non-linear transformation [28]. Mean ^18^F-Florbetapir uptake was measured across frontal, lateral parietal, lateral temporal and anterior/posterior cingulate. Then, the ^18^F-Florbetapir standardized uptake value ratio (SUVR) map was extracted using the whole cerebellum as reference region [29]. The PET scans of 5 participants were not suitable for ^18^F-Florbetapir quantification and were excluded of the quantitative analyses.

### 2.4 Statistical analysis

We performed receiver operating characteristic (ROC) analysis for Aβ1-42, Aβ1-40, tTau, pTau and the ratios Aβ1-42/Aβ1-40, Aβ1-42/tTau and Aβ1-42/pTau to calculate areas under the curve (AUC) with 95% confidence intervals (DeLong). We compared ROC curves applying a two-sided bootstrapping method with 2000 replications. For biomarkers and ratios that showed AUC higher than 0.70, we determined positive percent agreement (PPA or sensitivity) and negative percent agreement (NPA or specificity) and calculated optimal cutoffs maximizing their Youden J index (PPA + NPA - 1). We calculated the overall percent agreement (OPA) between CSF biomarker cutoffs and the amyloid PET visual interpretation as the sum of participants classified as "positive" or as "negative" by both modalities over the total number of participants. We also analyzed the correlation of each biomarker with global amyloid accumulation by fitting quadratic linear models and calculated the agreement of CSF biomarkers cutoffs with the PET scans quantification status applying a previously described SUVR cutoff of 1.11 [24]. Level of significance was set at α=0.05. We used Analyse-it^®^ statistical software for the selection of optimal cutoffs and packages “car”, “pROC”, “grid” and “ggplot2” as implemented in R statistical software (v 3.3.2) for plots and statistical analyses.

## 3. Results

### 3.1 Study participants

We included 94 participants in the study. **Table 1** summarizes demographic characteristics and biomarker results in the overall study population and according to the visual interpretation of amyloid PET scans as amyloid-positive (63%) or amyloid-negative (37%). There were no differences in age or sex between both groups. As expected, the amyloid-positive group had a higher proportion of *APOE*ε*4* carriers compared to the amyloid-negative group (52% and 20%, respectively; p=0.004).

**Table 1.** Clinical and demographic characteristics of all participants and based on visual amyloid PET status

|  | **All participants** | **Amyloid positive** | **Amyloid negative** | **P value** |
| --- | --- | --- | --- | --- |
| n (%) | 94 (100%) | 59 (63%) | 35 (37%) | - |
| Age, years | 73.0 (7.6) | 73.5 (7) | 72.1 (8.5) | 0.405* |
| Sex, Female/Male (% Female) | 50/44 (53%) | 32/27 (54%) | 18/17 (51%) | 0.960† |
| APOEε4 +/- (% +) | 56/37 (60%) | 30/28 (52%) | 7/28 (20%) | 0.004† |
| MMSE score | 24.7 (4.1) | 24.1 (4.3) | 25.7 (3.6) | 0.052* |
| Time difference between amyloid PET and lumbar puncture, days | 152 (86) | 151 (78) | 156 (100) | 0.792* |
| **Clinical diagnosis, n (%)** |  |  |  | 0.067† |
| Cognitively normal | 6 (100%) | 3 (50%) | 3 (50%) | - |
| Mild cognitive impairment | 35 (100%) | 23 (66%) | 12 (34%) | - |
| AD dementia | 12 (100%) | 11 (92%) | 1 (8%) | - |
| Dementia with Lewy bodies | 30 (100%) | 18 (60%) | 12 (40%) | - |
| Frontotemporal dementia | 9 (100%) | 3 (33%) | 6 (67%) | - |
| Other diagnoses | 2 (100%) | 1 (50%) | 1 (50%) | - |
| **CSF biomarkers** |  |  |  |  |
| Aβ1-42, pg/ml | 745 (379) | 608 (213) | 985 (474) | <0.001* |
| Aβ1-40, pg/ml | 12252 (3944) | 12782 (3954) | 11360 (3816) | 0.089* |
| tTau, pg/ml | 566 (363) | 667 (375) | 397 (273) | <0.001* |
| pTau, pg/ml | 93 (71) | 114 (71) | 58 (58) | 0.001* |
| Aβ1-42/Aβ1-40 | 0.064 (0.028) | 0.049 (0.015) | 0.088 (0.027) | <0.001* |
| Aβ1-42/tTau | 1.89 (1.43) | 1.18 (0.80) | 3.09 (1.47) | <0.001* |
| Aβ1-42/pTau | 13.7 (11.7) | 7.4 (5.9) | 24.3 (11.4) | <0.001* |

Unless otherwise specified, results are presented as mean (standard deviation).
MMSE: Mini-Mental State Examination; AD: Alzheimer's disease; CSF: cerebrospinal fluid
P-values were calculated by comparing amyloid-positive and amyloid-negative participants using Welch two-sample t-test (\*) or Fisher's exact test (†)

### 3.2 Quantification of A**β**1-42, A**β**1-40, tTau and pTau concentrations on the LUMIPULSE G600II

We measured Aβ1-42, Aβ1-40, tTau and pTau levels simultaneously on the Lumipulse G System. Levels of biomarker quantifications in the overall study population ranged from 315 to 2280 pg/ml for Aβ1-42, 4585 to 25925 pg/ml for Aβ1-40, 141 to 1902 pg/ml for tTau, and 18 to 340 pg/ml for pTau.

The analyses were divided over three calibration runs on the LUMIPULSE G600II, and the calibration status was valid for all samples. In our study, mean inter-assay coefficients of variation for the Lumipulse controls were 2.8%, 2.9%, 2.2% and 5.5% for Aβ1-42, Aβ1-40, tTau and pTau, respectively (**Supplementary Figure 1**).

A different aliquot of most of the CSF samples included in this study had previously been analyzed using other immunoassays, and their results were available in our database. Although these historic results were obtained in the context of routine clinical assessment by using different batches, and therefore a side-to-side precision analysis could not be performed, we explored their correlation with the Lumipulse G quantifications. The Lumipulse G assays for Aβ1-42, tTau and pTau showed very high correlation with values previously measured with Fujirebio’s INNOTEST ELISA (Pearson’s r of 0.94, 0.95 and 0.95, respectively, all p<0.001). The Lumipulse G assay for Aβ1-40 showed moderate correlation with values measured with Merck-Millipore’s ELISA (Pearson’s r of 0.76, p<0.001, **Supplementary Figure 2**).

### 3.3 Agreement between ^18^F-Florbetapir visual status and CSF biomarkers

We performed ROC analysis to investigate the accuracy of each CSF biomarker to detect the visual status of ^18^F-Florbetapir amyloid scans.

As displayed in **Figure 1A**, of all four individual biomarkers, tTau and pTau had the highest accuracy and showed AUC of 0.80 (95% CI 0.70-0.89, p<0.001) and 0.84 (95% CI 0.75-0.93, p<0.001), respectively. Aβ1-42 had fair accuracy with an AUC of 0.76 (95% CI 0.65-0.86, p<0.001) and Aβ1-40 alone was not useful for the detection of the visual status of amyloid scans (AUC 0.59; 95% CI 0.47-0.71, p=0.134).

**Figure 1.**
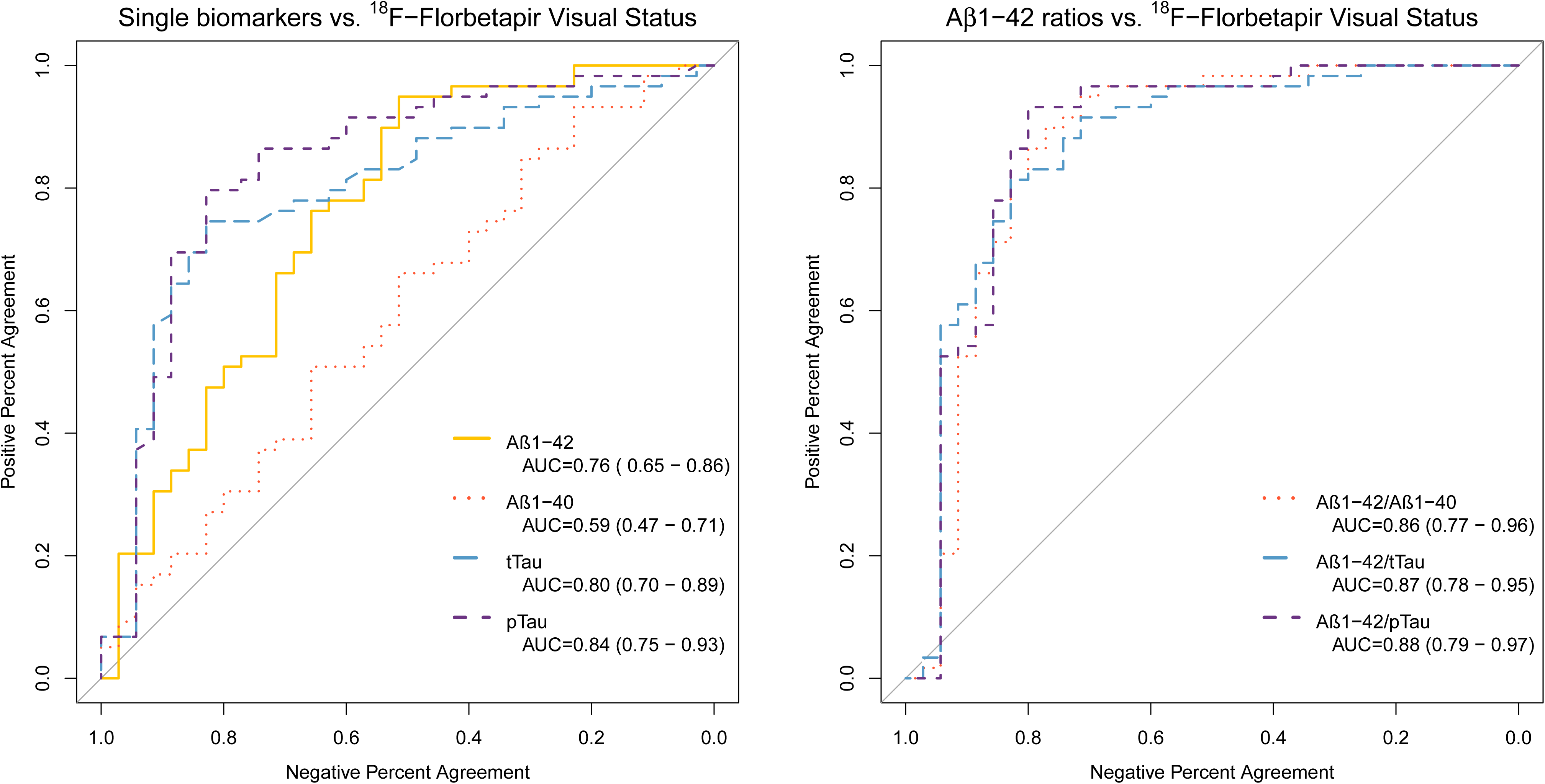
Receiver operating characteristic analysis of individual (A) and combined (B) CSF biomarkers’ diagnostic utility to detect amyloid visual status. AUC: Area under the curve.

**Figure 1B** shows that the combination of Aβ1-42 with a second analyte resulted in significant increases of accuracy. Aβ1-42/Aβ1-40 obtained an AUC of 0.86 (95% CI 0.77-0.96, p<0.001), significantly higher than that of Aβ1-42 alone (D=-2.5; p=0.01) or Aβ1-40 alone (D=-4.0; p<0.001). Aβ1-42/tTau had an AUC of 0.87 (95% CI 0.78-0.95, p<0.001), significantly higher compared to that of tTau (D=-2.2; p=0.03) and, Aβ1-42/pTau had and AUC of 0.88 (95% CI 0.79-0.97, p<0.001), higher than that of pTau alone (D=-1.9; p=0.05). Combining a third biomarker in the ratio did not improve its accuracy (data not shown).

### 3.4 CSF biomarker cutoffs based on visual interpretation of amyloid status

For those biomarkers and ratios that showed AUC higher than 0.70, we used ROC analysis to obtain PPA, NPA and OPA for all possible cutoffs. We chose optimal cutoffs for each biomarker and ratio by maximizing their Youden J index (PPA + NPA - 1). This approach is equivalent to maximize accuracy for a disease prevalence of 50% [30]. As displayed in **Figure 2**, in the case of single biomarkers, Aβ1-42, tTau and pTau, the selection was based on clear Youden peaks at 916 pg/ml, 456 pg/ml and 63 pg/ml, respectively. For the ratios Aβ1-42/Aβ1-40, Aβ1-42/tTau and Aβ1-42/pTau, plots showed *plateau* stages indicating that a wide range of cutoffs yielded similar Youden J indices. Best cutoffs for ratios were 0.062 for Aβ1-42/Aβ1-40, 1.62 for Aβ1-42/tTau, and 15.1 for Aβ1-42/pTau.

**Figure 2.**
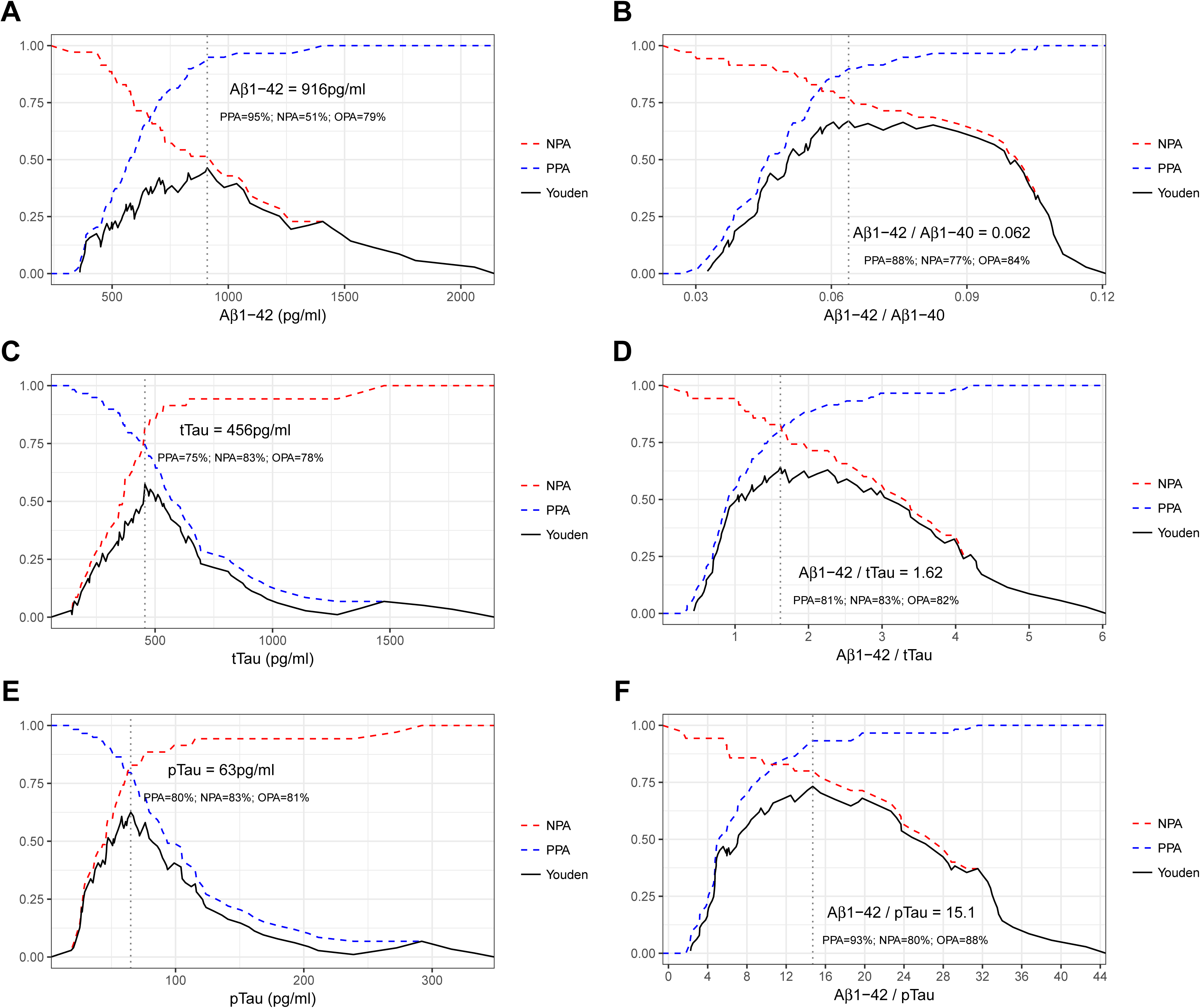
Accuracy of all possible cutoff levels of individual (A, C, E) and combined (B, D, F) CSF biomarkers. Only those biomarkers that yielded areas under the curve above 0.70 and their ratios with Aβ1-42 are shown. Vertical dotted lines indicate cutoffs with maximum Youden J index. PPA: Positive Percent Agreement; NPA: Negative Percent Agreement; OPA: Overall Percent Agreement

**Figure 3 A-C** displays the agreement between visual status of ^18^F-Florbetapir and CSF biomarker cutoffs that were optimal in our study for each analyte and ratio. For Aβ1-42, tTau and pTau, the OPA values between visual status and CSF biomarkers status were 79%, 78% and 81%, respectively. The ratio of Aβ1-42 with Aβ1-40, tTau and pTau increased the OPA to 84%, 82% and 88%, respectively.

**Figure 3.**
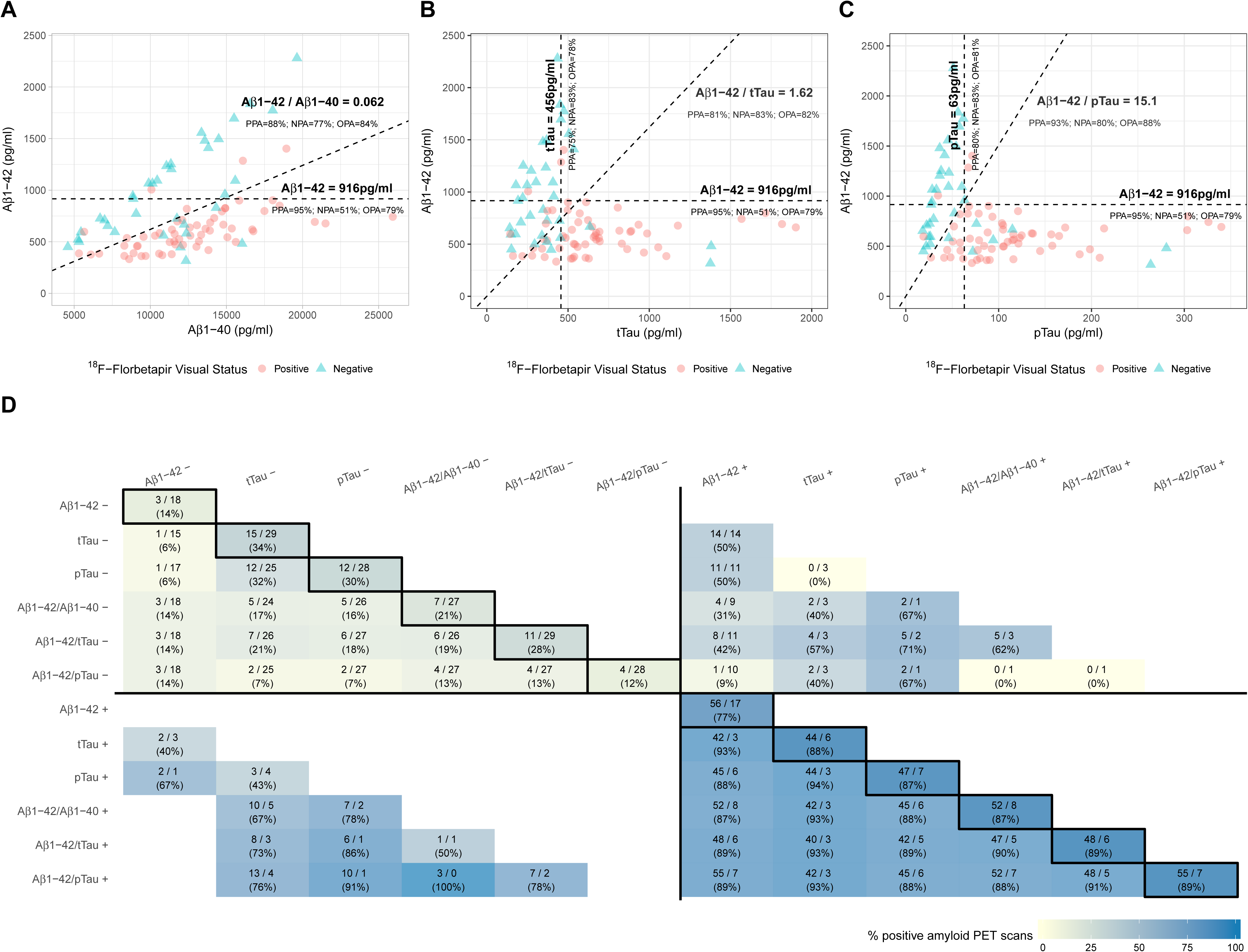
Agreement of visual amyloid status with single and combined CSF biomarkers. Panels A, B and C display scatterplots of CSF biomarker levels. Dashed lines indicate cutoffs that yielded maximum Youden J Index in the receiver operating characteristic analysis for each biomarker or ratio. Panel D illustrates the number of positive and negative amyloid PET scans for each CSF biomarker or ratio status (bordered cells, in the diagonal) and when a combination of two CSF biomarkers or ratios is considered (non-bordered cells). Values are represented as positive scans / negative scans (percent of positive scans). PPA: Positive Percent Agreement; NPA: Negative Percent Agreement; OPA: Overall Percent Agreement

### 3.5 Markers of amyloid and importance of assessing a second biomarker to predict the visual amyloid status

The proportion of positive amyloid scans varied significantly within each CSF biomarker status when a second biomarker or ratio was taken into account. **Figure 3D** shows the proportion of positive amyloid scans within each combination of two CSF biomarkers or ratios, and illustrates the importance of considering a second biomarker. Of all participants with low CSF levels of Aβ1-42 (below the cutoff of 916 pg/ml), regardless of the Aβ1-42/Aβ1-40 ratio status, 77% (56 out of 73) had a positive amyloid scan. This proportion increased to 87% (52 out of 60) within this group when the Aβ1-42/Aβ1-40 ratio was known to be also low (below 0.062) but decreased to 31% (4 out of 13) when this ratio was high (above 0.062). In the group of participants with high CSF levels of Aβ1-42 (above the cutoff of 916 pg/ml), the impact of considering the Aβ1-42/Aβ1-40 ratio status had no effect, as in all participants within this group the Aβ1-42/Aβ1-40 ratio was also high. These results highlight the importance of using the Aβ1-42/Aβ1-40 ratio over Aβ1-42 alone in the assessment of brain amyloidosis, especially in patients with low CSF levels of Aβ1-42.

### 3.6 Agreement of CSF cutoffs with amyloid quantification

We next processed amyloid PET scans to obtain quantification values of amyloid deposition. In our study, the previously validated SUVR value of 1.11[29], showed 83% PPA, 76% NPA, and 81% OPA with visual classification. As displayed in **Supplementary Figure 3**, scans that were divergently classified as “negative” or “positive” by one of the three raters showed intermediate SUVR values compared to scans that were unanimously classified.

As seen in **Figure 4**, the agreement of CSF cutoffs with amyloid PET quantification was similar to that with visual classification for all individual biomarkers and ratios. We studied the correlation between each CSF biomarker and global amyloid accumulation by fitting quadratic linear models. In these models, the adjusted coefficients of determination (*R*^*2*^) for individual biomarkers were 0.22 for Aβ1-42 (p<0.001), 0.03 for Aβ1-40 (p=0.06), 0.27 for tTau (p<0.001) and 0.38 for pTau (p<0.001). The adjusted coefficients of determination were higher for all ratios compared to individual biomarkers. The combination of Aβ1-42 with Aβ1-40 increased *R*^*2*^ value to 0.44 (p<0.001), indicating that this ratio reflects the amyloid deposition better than Aβ1-42 alone. Stratified analysis by ^18^F-Florbetapir visual status showed lower *R*^*2*^ values for all biomarkers, which suggests that the correlation observed between CSF biomarkers and amyloid PET quantification is partially mediated by an amyloid-status or diagnostic effect. In this stratified analysis, the highest correlation of SUVR values was found with the Aβ1-42/Aβ1-40 ratio in the amyloid-negative group (*R*^*2*^=0.42; p<0.001).

**Figure 4.**
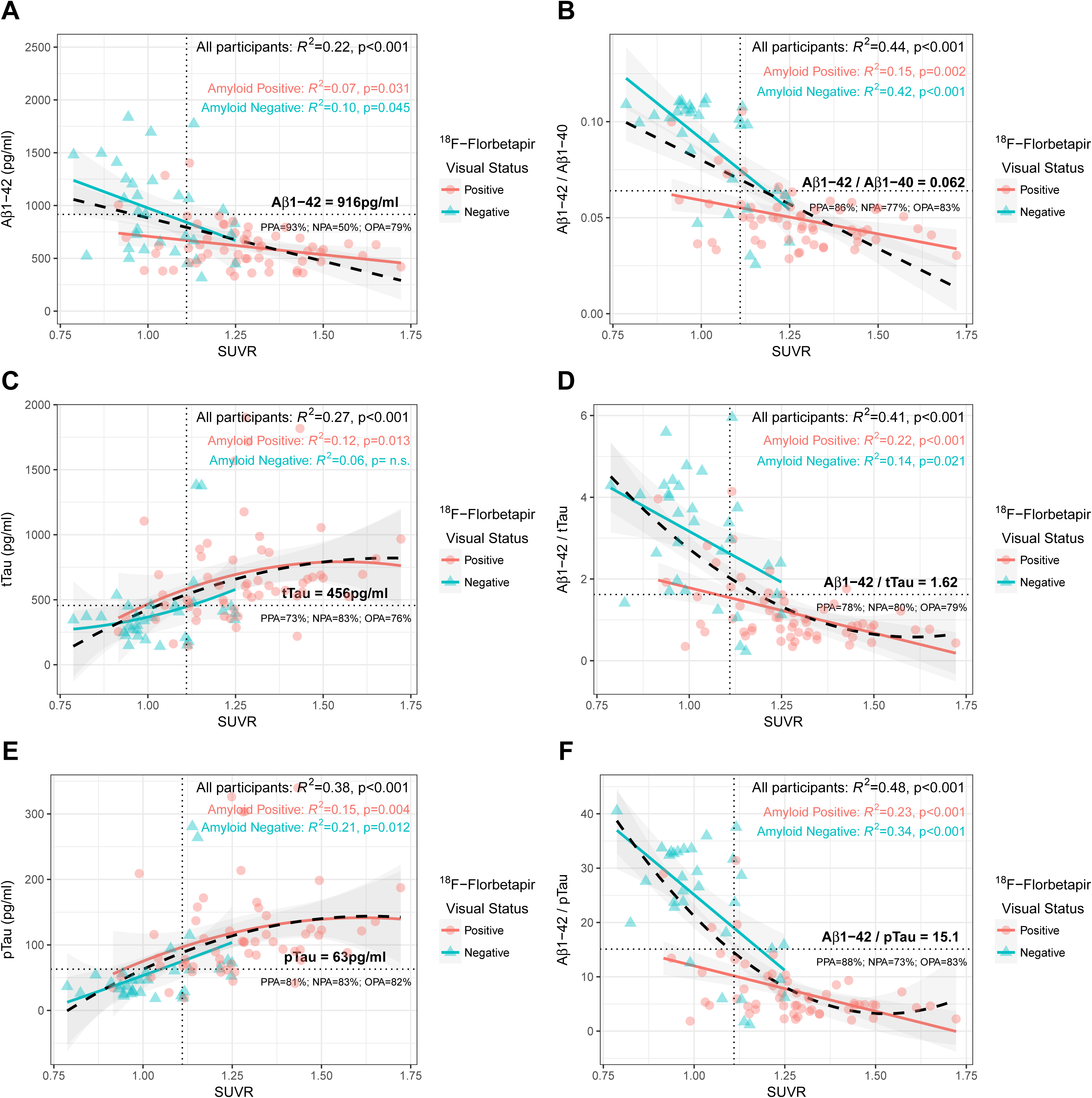
Scatterplots and correlations of amyloid quantification values with individual biomarkers (A, C, E) and ratios (B, D, F) Correlation between SUVR values and CSF biomarkers was assessed by fitting quadratic linear models for all participants (black) and after stratifying by visual amyloid status (red and green). Shaded areas indicate 95% confidence intervals. Dashed vertical lines indicate the SUVR cutoff of 1.11 as in Landau et al. Horizontal lines correspond to cutoffs for each CSF biomarker and ratio. PPA: Positive Percent Agreement; NPA: Negative Percent Agreement; OPA: Overall Percent Agreement; SUVR: Standardized Uptake Value Ratio

## 4. Discussion

In our study, we determined cutoffs for four CSF biomarkers for AD (Aβ1-42, Aβ1-40, tTau and pTau) and their ratios measured on the fully automated LUMIPULSE G600II platform to optimize their concordance with ^18^F-Florbetapir PET. We calibrated Aβ1-42 levels to certified reference material, recently developed to harmonize immunoassays across different platforms, and found that the ratios Aβ1-42/Aβ1-40, Aβ1-42/tTau and Aβ1-42/pTau had better diagnostic agreement with visual assessment of amyloid scans than single biomarkers. As a marker of amyloid pathology, the Aβ1-42/Aβ1-40 ratio had higher agreement with amyloid PET visual status and showed better correlation with amyloid load quantification compared to Aβ1-42 alone.

The agreement between amyloid imaging and AD CSF biomarkers has previously been studied by using other automated immunoassays [10,11,17]. Our results are in line with previous studies showing a good overall agreement between amyloid imaging and AD CSF biomarkers, higher for ratios than for single analytes [10,11]. However, specific cutoff points for CSF biomarkers differ between these studies, and several methodological differences can explain these discrepancies. First, pre-analytical conditions, such as the type of collection and storage tubes, are different between studies, and these factors are known to have a great impact on the absolute values of CSF biomarkers, especially for Aβ1-42 [31,32]. Second, some analytical particularities for each immunoassay and platform used in these studies (specificity of the antibodies, time of incubation) result in diverse CSF biomarker measures. Calibration of all automated platforms to certified reference material, currently underway, will minimize this issue in the future. Likewise, differences in the affinity of PET radiotracers (^11^C-Pittsburg compound B, ^18^F-Flutemetamol or ^18^F-Florbetapir) can lead to disparities in the selection of cutoffs. Third, the definition of the study population can have an impact on the measures of diagnostic accuracy and the determination of cutoffs, and the composition of the populations was not the same across studies. Schindler et al. analyzed data from community-dwelling volunteers [11], whereas Janelidze et al. obtained their results from patients with mild cognitive impairment and subjective cognitive decline from the BioFINDER cohort [17]. Hansson et al. studied CSF of participants from ADNI and BioFINDER cohorts, that included cognitively normal volunteers, patients with mild cognitive impairment and patients with AD dementia [10]. In our study, we additionally included patients with other dementias or neurodegenerative diseases, which might reflect more realistically the application of biomarkers in daily clinical practice.

Besides these technical particularities, the definition of what is an optimal cutoff should finally depend on the aim of the test. As in a number of other studies, the cutoffs in our study were selected by maximization of Youden J index. This approach balances sensitivity and specificity and is equivalent to maximize accuracy for a pre-test disease prevalence of 50% [30]. However, other strategies might be useful in certain clinical scenarios. For instance, for screening purposes, it might be helpful to apply cutoffs with high sensitivity, even when their specificity is lower. For patients with clinically challenging diagnoses, however, high specificity might be preferable. Other possible approaches include the sequential application of biomarker cutoffs [33].

The LUMIPULSE G600II has incorporated the possibility of measuring CSF levels of Aβ1-40. In previous studies, and ours, this biomarker alone was not useful for the detection of brain amyloid [11,14,18], but its use in combination with Aβ1-42 increased significantly the accuracy of Aβ1-42 alone. Both Aβ1-42 and the Aβ1-42/Aβ1-40 ratio are included in the “A” category of the ATN classification system together with amyloid PET, but in line with other studies [14,18,34,35], we found that the Aβ1-42/Aβ1-40 ratio had better agreement with visual amyloid status and higher correlation with brain amyloid quantification. Our results also suggest that the use of the Aβ1-42/Aβ1-40 ratio could be crucial to compensate individual differences in amyloid precursor protein processing that otherwise might lead to false positive or false negative Aβ1-42 CSF levels. In our study, this was particularly relevant for participants with low CSF levels of Aβ1-42 levels. This information adds to the fact that using the Aβ1-42/Aβ1-40 ratio has proven to partially mitigate the effect of some pre-analytical confounders that have been described to alter the results of amyloid levels [36,37] (Delaby et al., under review). Altogether, our data suggest that the use of the Aβ1-42/Aβ1-40 ratio would be more reliable in clinical practice than Aβ1-42 alone as a marker of amyloidosis and that this combination should be used in routine, being particularly relevant in cases with low Aβ1-42 levels.

The main strength of our study is that four AD CSF biomarkers (Aβ1-42, Aβ1-40, tTau and pTau) measured simultaneously with the fully automated Lumipulse G System were compared for the first time to ^18^F-Florbetapir PET to calculate amyloid-based cutoffs. In addition, this is, to our knowledge, the first study to present Aβ1-42 levels that have been standardized to certified reference material, recently developed to harmonize immunoassays across different platforms. The standardized values that we present will make our study more easily comparable to future studies. Moreover, to avoid possible sources of variability, we followed homogeneous CSF pre-analytical and analytical procedures and used the same batch of reagents for all measurements. Also, the inclusion of participants with neurodegenerative diseases outside the AD spectrum provides a more realistic application of biomarkers in daily clinical practice.

However, we acknowledge that our study has also some limitations. We did not test the effect that deviations from our pre-analytical protocol would have on the exact final cutoffs, and therefore, the cutoffs that we report should be taken cautiously under other operating procedures. Additionally, only very few participants had additional Tau imaging and/or ^18^F-Fluorodeoxyglucose-PET, and therefore, we could not compare the agreement of CSF pTau and tTau with molecular imaging markers of the “T” and the “N” categories of the ATN classification system. Likewise, as participants in this study are part of a living cohort, neuropathological confirmation is not available at this moment.

In this study, we found that the Aβ1-42 ratios to Aβ1-40, tTau and pTau in CSF show a good agreement with amyloid visual status and that the Aβ1-42/Aβ1-40 ratio had better correlation with the amount of amyloid burden compared to Aβ1-42 alone. The understanding of the agreement between CSF biomarkers and amyloid imaging is crucial to identify situations in which these two modalities might not be interchangeable. This information has to be taken into consideration both in the diagnostic assessment in clinical practice and in the selection of participants in clinical trials.

## Supporting information

## Acknowledgements

We are grateful to all participants in the study and to all the members of the clinical team that were involved in the selection and assessment of participants.

This study was supported by CIBERNED and Instituto de Salud Carlos III PI13/01532 and PI16/01825 to R.B.; PI14/01561 and PI17/01895 to A.L; PI14/01126 and PI17/01019 to J.F., jointly funded by Fondo Europeo de Desarrollo Regional (FEDER), Unión Europea, “Una manera de hacer Europa”. This work was also supported by Generalitat de Catalunya (2017-SGR-547, SLT006/17/125 to D.A., SLT006/17/95 to E.V. and SLT006/17/119 to J.F.) and “Marató TV3” foundation grants 20141210 to J.F., 20161431 to R.B. and 20142610 to A.L.

All kits and reagents to perform cerebrospinal fluid analyses in this study were provided by Fujirebio Europe.

