## Supplementary material for "Agreement between ^18^F-Florbetapir PET imaging and cerebrospinal fluid Aβ1-42, Aβ1-40, tTau and pTau measured on the LUMIPULSE G fully automated platform"

**Supplementary Figure 1 - Inter-assay coefficients of variation calculated from internal quality controls with low, medium and high concentration of Aβ1-42 (A), Aβ1-40 (B), tTau (C) and pTau (D)**

CV: Coefficient of Variation

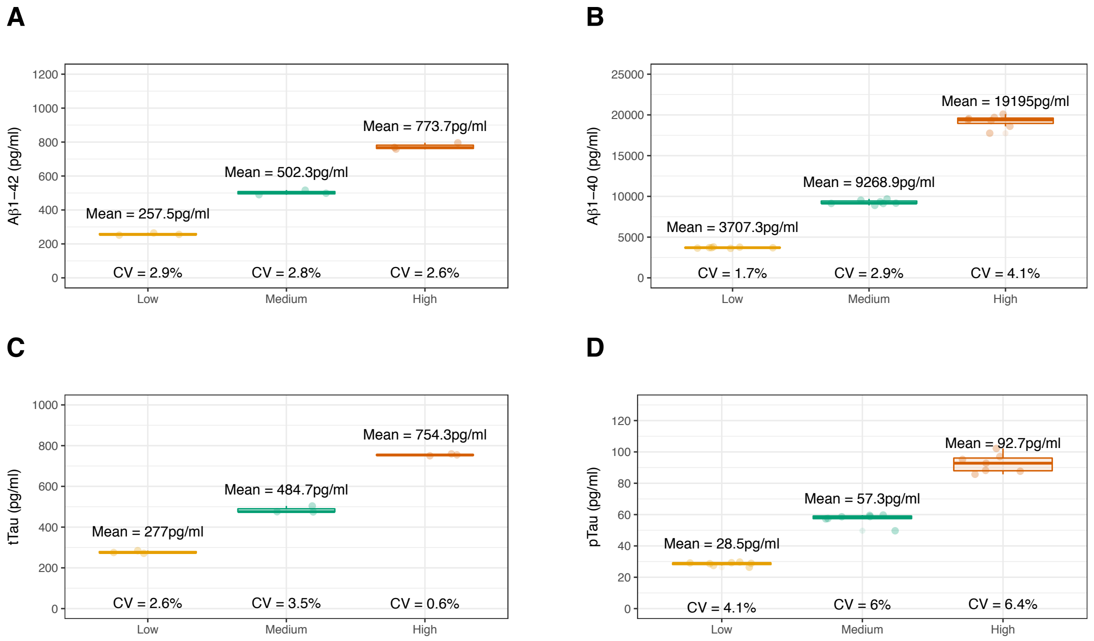

### Supplementary Figure 2 – Correlation between Lumipulse G analysis and historic ELISA values determined in the same samples

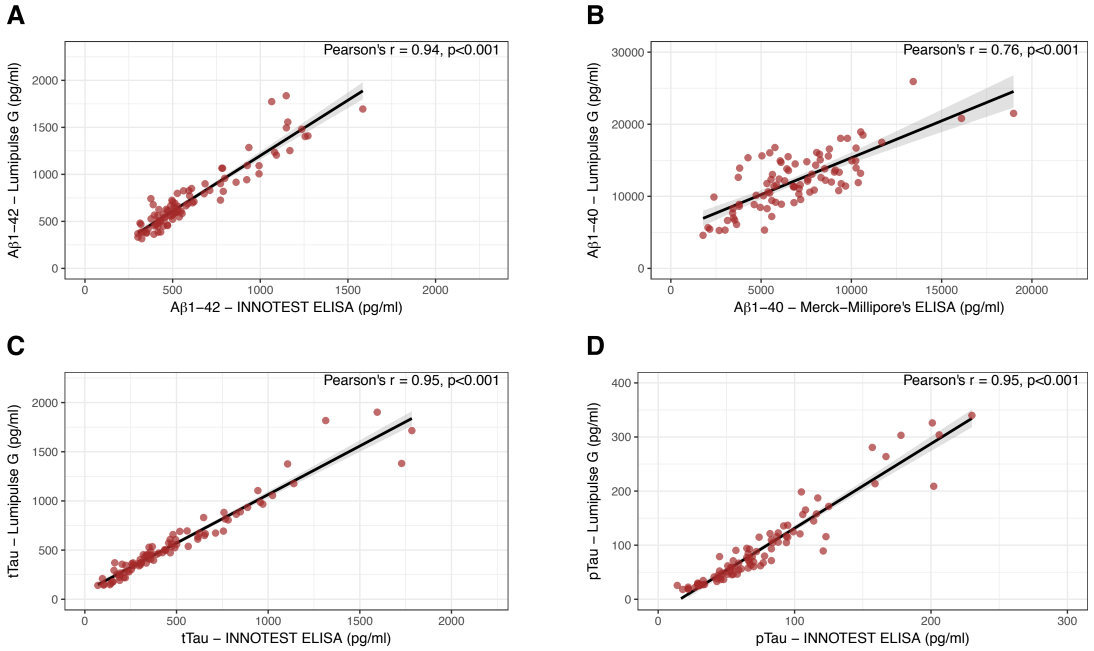

**Supplementary Figure 3 – Agreement between raters' visual classification and amyloid quantification**

OPA: Overall Percent Agreement; SUVR: Standardized Uptake Value Ratio

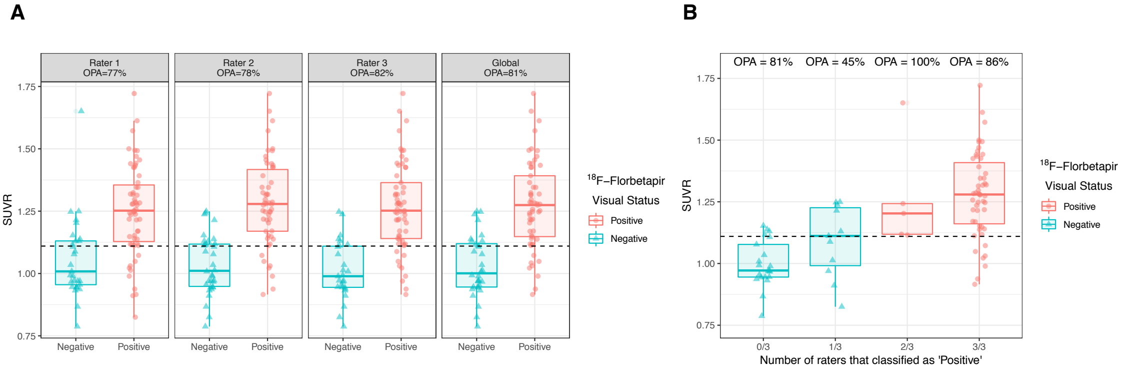
